# GWAShub: A Web-based Resource to Democratize Genome-Wide Association Studies in Crop Plants

**DOI:** 10.1101/2022.02.03.479034

**Authors:** Anurag Daware, Rishi Srivastava, Durdam Das, Naveen Malik, Akhilesh K. Tyagi, Swarup K. Parida

## Abstract

Genome-wide association study (GWAS) is a popular approach for linking natural genetic variation with phenotype variation and thus is central to crop quantitative genetics. The ever-increasing wealth of publicly available genomic sequence information for crop plants presents an unprecedented opportunity for utilizing GWAS for the identification of genes governing a plethora of agronomic traits. However, the lack of technical expertise and computational infrastructure is still hindering the ability of plant breeders to conduct GWAS in a self-reliant manner. Here, we present a GWAShub web server that provides a user-friendly interface for performing comprehensive GWAS and post-GWAS in crop plants utilizing publicly available genomic sequence variation data, comprehensive annotation data and diverse computational tools. The utility of GWAS-hub was further demonstrated by conducting large-scale GWAS for flowering/maturity time traits in chickpea. This analysis identified three different flowering/maturity time associated genes, all encoding different histone methyltransferases. Thus, epigenetic regulation is identified as vital mechanism regulating flowering time and maturity duration in chickpea. Finally, we hope GWAShub (www.gwashub.com) will enable resource-scarce researchers to join the GWAS revolution fueled by advancements in next-generation sequencing and computational genomics

## Introduction

Since its inception in maize, GWAS has become a widely popular and successful tool for dissecting the genetic basis of complex quantitative traits in crop plants (**Thornsberry et al., 2001**). The popularity and success of GWAS for trait mapping in crop plants can be primarily attributed to the lack of need for the generation of experimental mapping populations and higher mapping resolution compared to other quantitative trait loci (QTL) mapping approaches. Additionally, the crop diversity panels once genotyped, can be maintained indefinitely and can be repeatedly phenotyped for any number of phenotypes under multiple environments/agro-climatic conditions. This makes GWAS in crop plants a highly robust method for rapid genetic dissection of a highly complex trait in a time and cost-efficient manner. So far, GWAS has been successfully implemented in almost all major crop plants like maize, rice, wheat, soybean, chickpea, tomato (**Huang et al., 2010; Huang & Han, 2014; Sauvage et al., 2014; Xiao et al., 2017; Yano et al., 2016; Basu et al. 2019)**, etc. GWAS studies in these crop species led to the identification of many important genes/alleles regulating diverse agronomic traits including plant growth/developmental and yield-related traits (**Thornsberry et al., 2001; Leiboff et al., 2015; Si et al., 2016; Romero Navarro et al., 2017; Prince et al., 2019; Yano et al., 2019; Tao et al., 2020**), biotic stress tolerance (**Dimkpa et al., 2016; Raboin et al., 2016; Wen et al., 2019; Yates et al., 2019**), abiotic stress tolerance (**Albert et al., 2016; Melandri et al., 2020**), nutritional quality traits (**Angelovici et al., 2017; Misra et al., 2019**), nutrient-use-efficiency (**Tang et al., 2019; Yang et al., 2018**), etc. The outcome of GWAS research has been reviewed recently (Huang and Han 2014; Wang et al. 2020).

In the last decade, the rapidly declining cost of next-generation sequencing (NGS) and other high-throughput genotyping technologies has made available publicly accessible sequence variation data (at least raw sequence data) for a large number of accessions for many major and minor crop species such as maize, rice, soybean, chickpea, pegionpea, tomato, etc. This includes sequence variation data generated by large International Consortiums (**Huang et al., 2010; Lin et al., 2014; Zhou et al., 2015; McCouch et al., 2016; Bukowski et al., 2018; Wang et al., 2018 Torkamaneh et al., 2019; Varshney et al., 2019**) and those generated by individual efforts (**3K RGP, 2014; Bukowski et al. 2017, He et al 2019; Varshney et al., 2021**). Apart from the genomic sequence variation data, multiple open-source computational programs for performing and visualizing GWAS-related relevant information have also been developed further encouraging trait association study in crop plants (**Lipka et al., 2012; Chang et al., 2015; Huang et al., 2019**). Despite the availability of numerous publicly accessible sequence variation datasets and advanced open-source computational programs, the GWAS utilizing these big datasets still remains scarce in the scientific literature. This suggests that the wealth of enormous genomic information available in the public domain for multiple crop plants has remained largely untapped for their genetic improvement. The unavailability of high-end computational infrastructure and lack of advanced programming skills required to acquire, organize and utilize this large-scale crop genomic data for GWAS seems to be the dominant cause hindering the use of aforementioned sequence variation data by researchers routinely working with these crop species for genomics-assisted crop improvement.

Therefore, a web-based, user-friendly, computational platform that can enable plant researchers to overcome these aforesaid hurdles in utilizing publicly accessible huge genomic information for performing GWAS for complex trait dissection will be highly beneficial. Although some of the previously available web resources such as easyGWAS (**Grimm et al., 2017**), Matapax (**Childs et al., 2012**), GWAAP (**Seren et al., 2013**) provide open-source platforms to conduct GWAS, they lack few important features which limit their wider applicability as well as utility for conducting GWAS in crop plants. Primarily, most of these web-resources focus on the model plant *Arabidopsis*, i.e., either they do not provide the conducive interface/option to analyze data from other plant species except *Arabidopsis* (e.g. Matapax and GWAAP), or a user needs to provide the genotype along with phenotype (e.g. easyGWAS) for trait association study. This makes it either impossible or extremely challenging to use these resources, for researchers routinely performing GWAS in plant species other than *Arabidopsis*. Besides, the existing GWAS servers provide a very limited number of algorithms and also none of these servers provide an option to utilize recently developed multi-locus-based GWAS algorithms which provide higher statistical power for the detection of genomic loci associated with complex quantitative traits. Further, none of these available resources provides extensive information on genome-wide haplotype blocks as well as detailed structural and functional annotation of GWAS-derived genomic loci associated with traits. Consequently, the existing servers lack the functionality to conduct post-GWAS analysis for the identification of causal trait-associated genes underlying detected GWAS loci. As drawing meaningful inference from GWAS results requires the integration of a diverse type of biological information, it becomes extremely challenging to conduct comprehensive GWAS using the aforementioned existing web servers.

Here, we introduce GWAShub (www.gwashub.com), a web-based platform that provides plant researchers with easy to use interface for performing comprehensive GWAS analysis by integrating publically available sequence variation data of diverse crops with user-generated private phenotype data. The GWAShub supports the implementation of eight diverse single-locus and multi-locus GWAS algorithms for diverse user applications. Further, it also enables exploration of pleiotropy at the genome-wide level by providing 2D as well as 3D landscape for simultaneous visualization of multi-trait Manhattan plots for GWAS. Finally, GWAShub provides extensive functionality for conducting post-GWAS analysis including exploration of local LD patterns, extensive annotation of the genomic intervals, and machine learning (ML)-based causal gene prioritization (**Figure 1**). The current version of GWAShub readily provides access to genomic variation datasets for four major crops i.e. rice, wheat, maize and chickpea with new datasets being constantly added to perform GWAS for quantitative dissection of the complex genetic architecture of agronomic traits and crop genetic enhancement.

**Figure 1.**
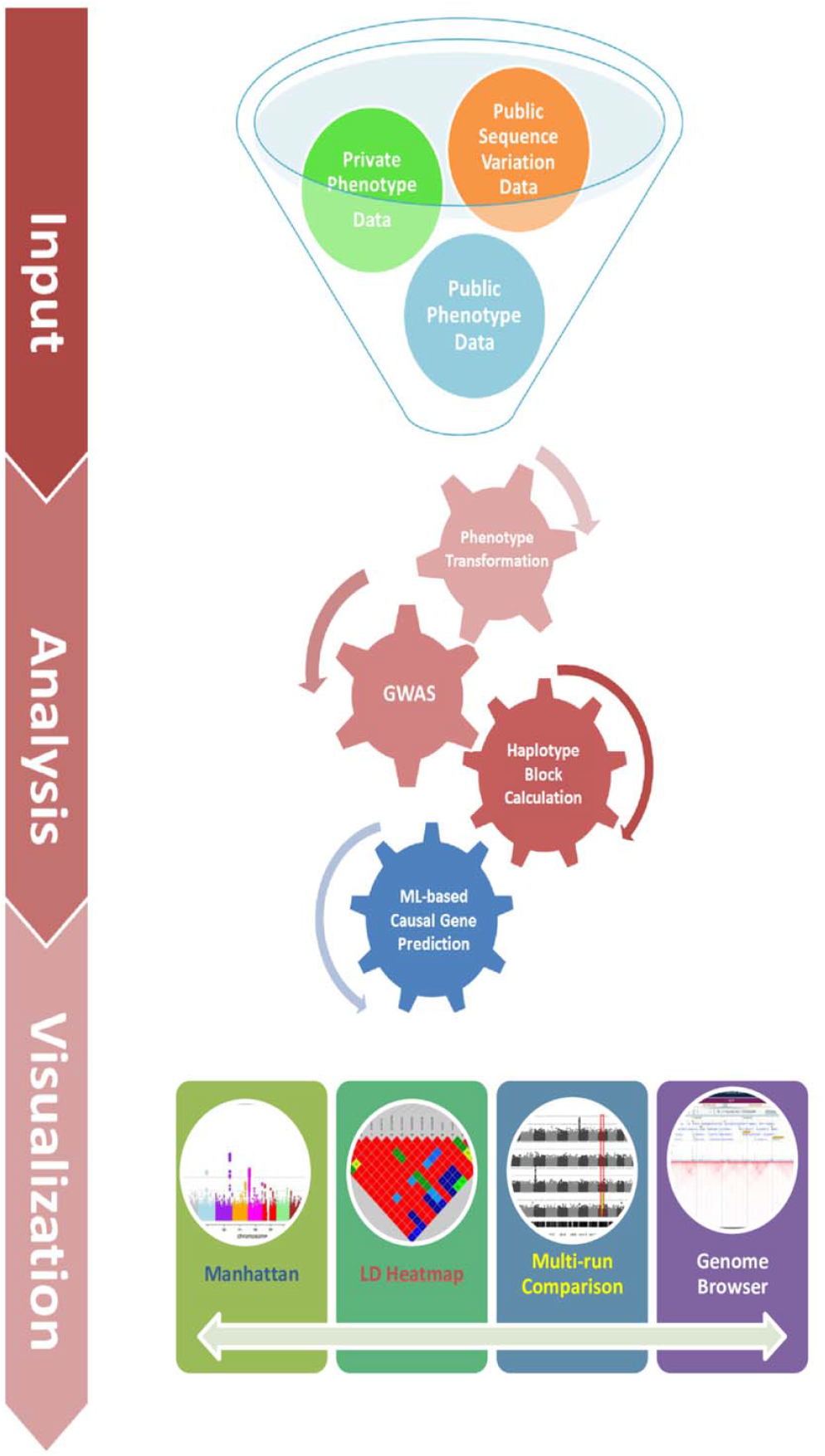
The diagrammatic overview of three major components of GWAShub webserver and its underlying functionalities. The three major components of GWAShub include Input, Analysis and Visualization.

### Data collection

In the last decade, rapid advancements in NGS technologies resulted in a significant increase in sequencing output and thus an exponential decline in sequencing cost. This inspired the global plant science community to undertake many large-scale genome sequencing efforts in almost all major crop plants. The high-quality sequence variation data for many crops are now available in the public domain. To include major cereal food crops, we retrieved genotype data (VCF format) for rice, wheat and maize from their respective databases (**Alexandrov et al., 2015; Bukowski et al., 2018; He et al., 2019**). Further, we also included chickpea as a representative legume food crop. As the variant data for chickpea is still not available publically, we downloaded the raw whole genome sequencing data of 402 diverse chickpea accessions from the SRA database (SRA: SRP096939; BioProject: PRJNA362278) (**Varshney et al. 2019)**. Subsequently, the sequence variants were identified as described by **Varshney et al. (2019)**. Further, SNP identified using whole-genome sequencing data of 3171 cultivated chickpea accessions (**Varshney et al. 2021**) was obtained from the CicerSeq database (https://cegresources.icrisat.org/cicerseq/). For the remaining three cereal crops, SNPs genotype data were obtained from their respective databases: Rice SNP-seek database (https://snp-seek.irri.org/) (**Alexandrov et al. 2015**), Maize HapMap (https://www.panzea.org/) (**Bukowski et al. 2018**), 1000 wheat exomes project (http://wheatgenomics.plantpath.ksu.edu/1000EC/) (**He et al. 2019**). The detailed information regarding reference assembly, number of accessions, number of SNPs and data source used in the present study for individual crops are provided in **Supplementary Table 1.**

## Methods

The GWAShub web-server was developed using Apache HTTP server (version 2.4.6) integrated with PHP (version 7.3.3) and MySQL (version 8.0.15) on a Centos 8 (Linux) operating system based server machine. R (version 3.6.2), JavaScript (version 1.8.0) and PHP were used to develop the back-end of the database while MySQL (version 8.0.15) was used to process the data at the back-end. CSS and HTML were used to make the template responsive. Multiple bash scripts were also integrated at the back-end of the database for multiple file handling, data conversion and manipulation.

Further, all the data transformation methods (except BoxCox) and a Shapiro-Wilk test for normality are implemented using core R functions, whereas, R library “Mass” was utilized to perform BoxCox transformation. Apart from this, all the eight GWAS algorithms implemented in GWAShub were adopted from GAPIT (Version 3.0), a popular R package routinely utilized for GWAS in crop plants (**Wang & Zhang, 2021**). For causal gene prioritization in postGWAS analysis, the QTG-finder (version 2.0) program was adopted with minor modifications (**Lin et al., 2020**). The QTG-finder (version 2.0) employs a supervised machine-learning (ML) approach and requires ML models trained using known causal genes, sequence variation data, functional annotation, gene essentiality and paralog copy number etc., for each of the respective crop species. For rice, the precomputed ML model available with QTG-finder (version 2.0) was utilized, while for the remaining three crop species we generated ML-models following **Lin et al. (2020)**. Further, the PheGWAS package which displays multi-trait Manhattan plots was used to display GWAS results in 3D landscape and in an interactive fashion. Similarly, the LDBlockShow program was utilized to generate LD heatmap displaying local LD patterns and haplotype block structures (**Dong et al., 2021**). Finally, the Jbrowse genome browser was integrated in GWAShub to visualize gene models as well as sequence variant data (**Buels et al., 2016**).

### PROGRAM DESCRIPTION

#### General description of the webserver

The GWAShub web-server is designed to conduct comprehensive GWAS and post-GWAS analysis in a user-friendly manner and currently harbors genotype datasets of four major food crops, namely rice, wheat, maize and chickpea (**Figure 2a**). GWAShub has three core components, viz. 1) Data input, 2) Analysis, and 3) Visualization. The data input component of GWAShub provides an interactive interface for the selection of pre-loaded genotype/phenotype datasets as well as for the upload of external user-generated phenotypes corresponding to the selected genotype dataset (**Figure 2b**). The analysis component has multiple independent interfaces for phenotype data transformation (**Figure 2c**), specifying GWAS run parameters (**Figure 2d**), and ML-based prioritization of causal genes (**Figure 2e**). Finally, the visualization component of GWAShub consists of diverse modules for detailed exploration of GWAS results (**Figure 3, 4**).

**Figure 2.**
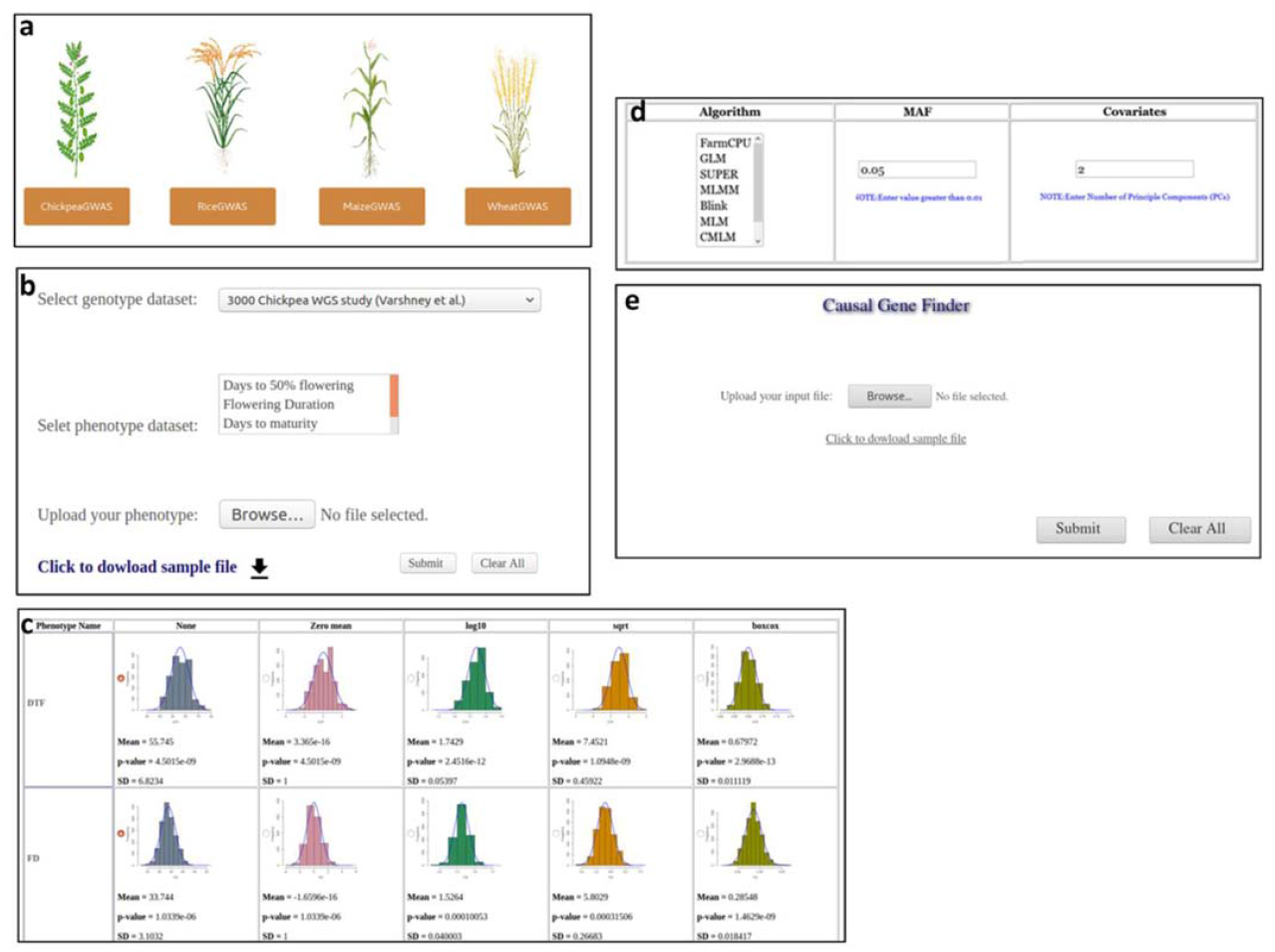
The data input and analysis components of GWAShub. **a)** Three major cereals and a legume crop species supported in the current version of GWShub. **b)** Interface for selection of genotypc/phenotype dataset and upload of user phenotype, **c)** Interface for selection of appropriate normalization method for user selected/ uploaded/phenotype data, **d)** Interface for setting optimal parameters for GWAS run. **e)** The post-GWAS causal gene finder interface where user can upload text file containing list candidate genes from multiple associated QTLs to prioritize the causal genes.

**Figure 3.**
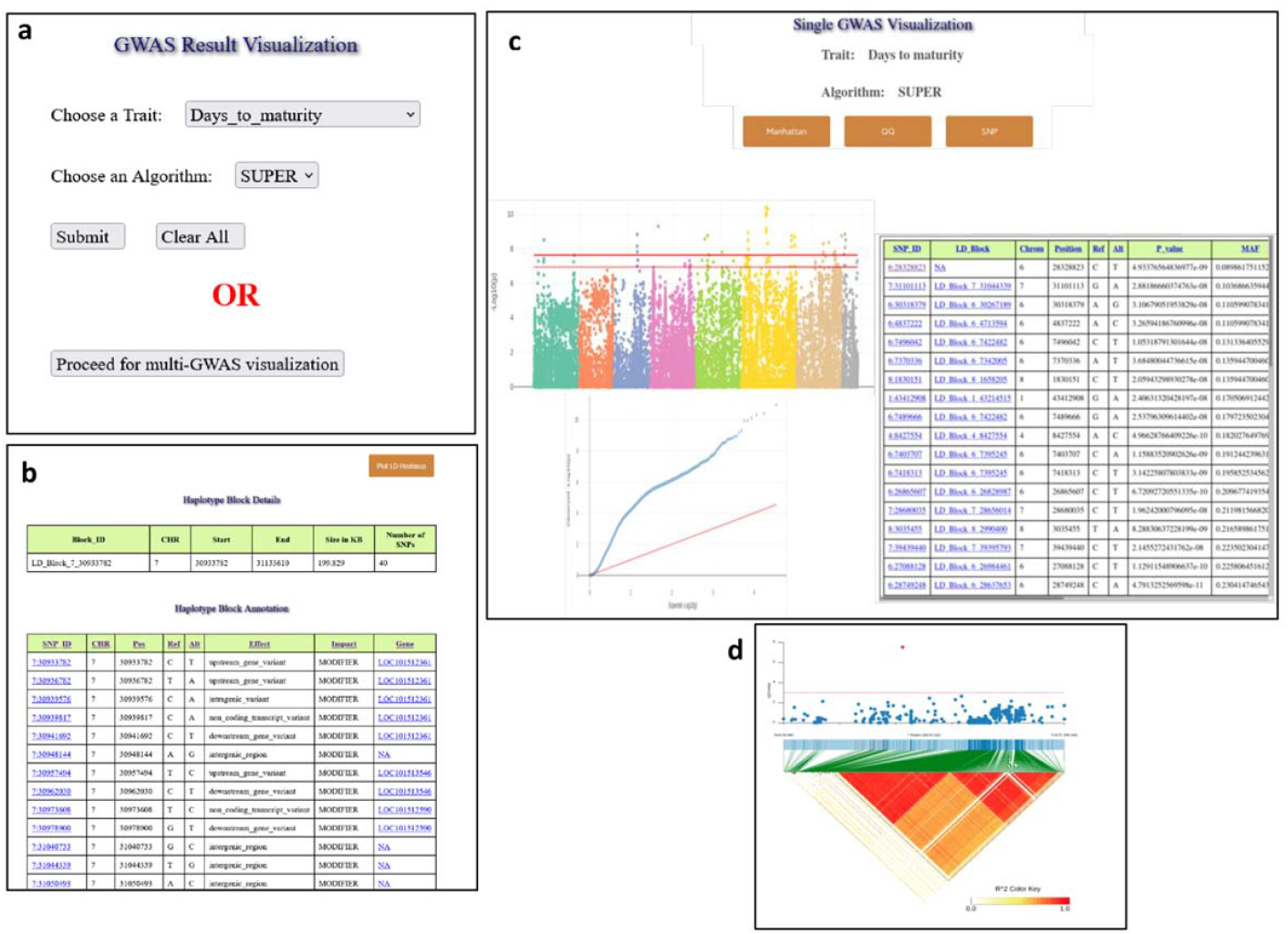
Visualization component of the GWAShuh web-server. **a)** Interface for selection of GWAS run i.e. trait and algorithm combination, **b)** Manhattan plot, QQ plot and association result table displayed for selected GWAS run, **c)** Table depicting summary of haplotype block and annotation of SNPs within the haplotype block, and **d)** LD-heat map displaying local LD for haplotype block along with its 200 kb upstream and downstream interval.

**Figure 4.**
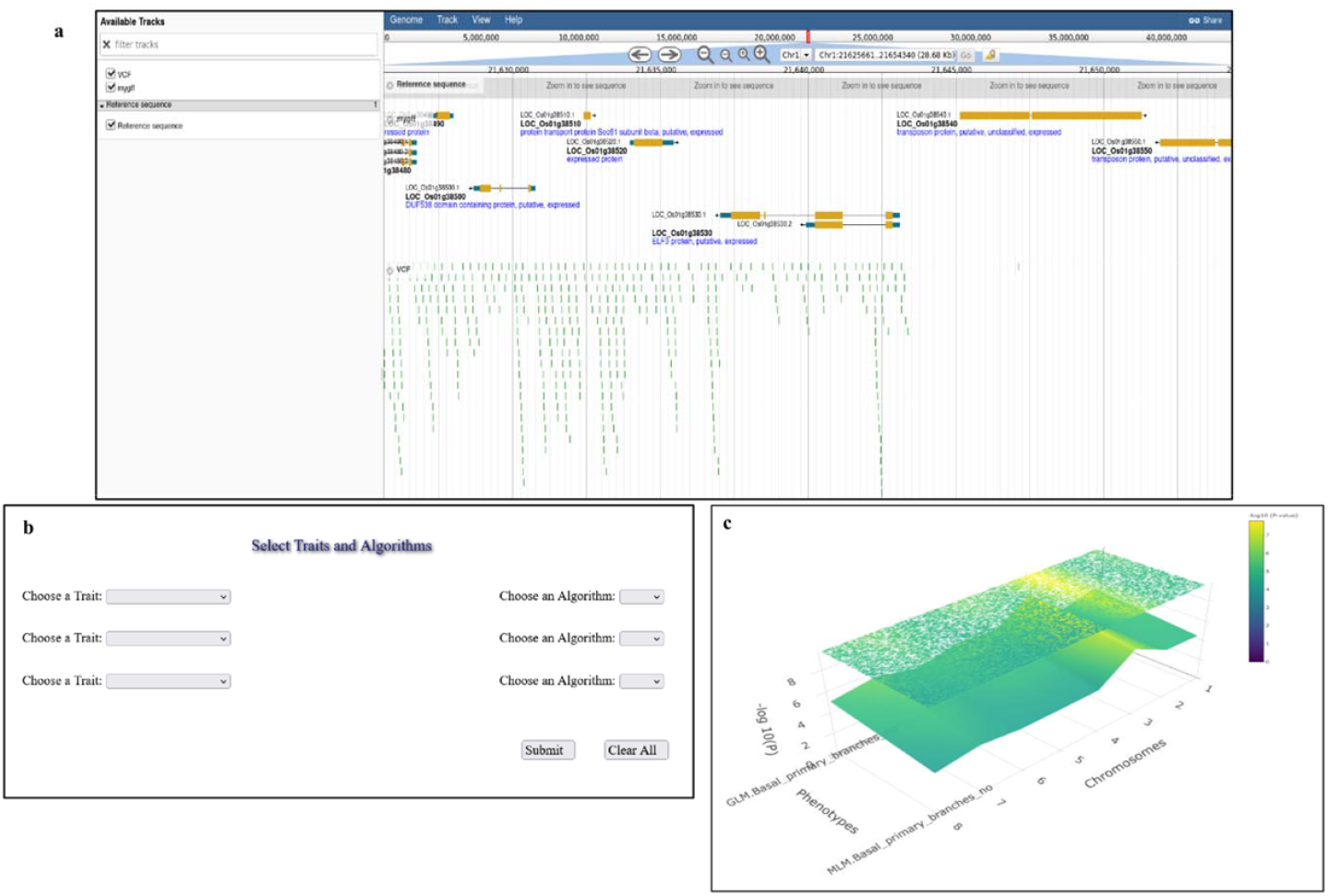
Visualization components of the GWAShub web server. **a)** Genome browser displaying gene models and sequence variants, **b)** Interface for selection of trait-algorithm combination for interactive 2D & 3D visualization of multiple traits/algorithms, and **c)** Interactive three dimensional plot of GWAS results for multiple traits/algorithms.

To initiate analysis on the GWAShub web-server, the user needs to click on the GWAShub button and select the crop of their interest (**Figure 2a**). The user then needs to select genotypes from the available genotype datasets and is further required to upload phenotypes corresponding to all or subset of accessions represented in the selected genotype dataset (**Figure 2b**). To improve the normality of phenotype, GWAShub has an option to select from four different types of transformation methods i.e. zero mean, log10, square-root, and Box-Cox. The information regarding phenotype distribution (histogram) and normality (Shapiro-Wilk test P-value) is displayed for each phenotype to assist with the selection of optimal transformation for each phenotype (**Figure 2c**). This is especially important as most statistical models used for GWAS are optimized for quantitative phenotypes with a normal distribution. After, the phenotype transformation, the next interface for GWAS parameters allows users to select one or more algorithms of their choice, minor allele frequency (MAF) and covariates (i.e. number of principal components) to be utilized for GWAS (**Figure 2d**). The current version of GWAShub provides eight different single/multi-locus GWAS algorithms which include generalized linear model (GLM) (**Wilson et al., 1999; Pritchard et al., 2000**), mixed linear model (MLM) (**Zhu et al., 2008**), compressed mixed linear model (CMLM) (**Zhang et al., 2010**), multiple loci mixed linear model (MLMM) (**Segura et al., 2012**), Settlement of MLM Under Progressively Exclusive Relationship (SUPER) (**Wang et al., 2014**), Fixed and random model Circulating Probability Unification (FarmCPU) (**Liu et al. 2012**), Bayesian-information and Linkage-disequilibrium Iteratively Nested Keyway (BLINK) (**Huang et al., 2019**) and multi-locus random-SNP-effect MLM (mrMLM) (**Wang et al., 2016**). Apart from GWAS, the current version of GWAShub provides utility for performing post-GWAS analysis aiming at the identification of causal genes underlying associated GWAS loci. The post-GWAS module accepts a list of candidate genes and ranks them based on their probability of being causal (**Figure 2e**). This module leverages a supervised machine learning algorithm for the prioritization of causal genes to rapidly identify the most likely major effect genes underlying each of the detected genomic loci.

After the successful completion of the GWAS run, the visualization component of GWAShub allows for further exploration of GWAS results. The results of each GWAS run can be visualized by selecting the appropriate trait-algorithm combination (**Figure 3a**). These results include the interactive Manhattan plots, QQ-plots and GWAS result table containing the SNP-ID, chromosome, haplotype block, P-value, MAF, false discovery rate (FDR)-adjusted P-value, variant annotation, predicted functional effect, etc., corresponding to top 50 trait-associated SNPs (with least P-values) (**Figure 3b**). The user can further explore the detected trait-associated loci in detail by clicking on SNP-IDs provided in the GWAS result table, which opens the corresponding positions (bp) of target loci in the genome browser (**Figure 3c**). The details of the haplotype block associated with SNP of interest and along with detailed annotation of SNPs within the haplotype block can be readily obtained by clicking on LD block IDs. Further, local LD patterns within and around the haplotype block can be visualized in the form of an LD heatmap by clicking on the Plot LD heatmap button (**Figure 3d**). This integrated LD heatmap image can also be retrieved by clicking on the download button provided on the top right corner of the image (**Figure 3d**). The GWAShub genome browser simultaneously displays multiple tracks including gene models and sequence variants. Further, the sequence variants are coded with different colors depicting the severity of the effect (low, moderate and high) caused by the variants (**Figure 4a**). This is vital for examining the GWAS-detected loci for genes with major effect mutations and therefore can delineate the potential candidate genes associated with traits of agronomic importance. Finally, GWAShub users can choose to simultaneously visualize and compare GWAS results for different trait-algorithm combinations by generating highly interactive 2D as well as 3D plots (**Figure 4b)**. Especially the 3D plot displaying results from multiple traits can enable efficient detection of pleiotropic loci regulating diverse agronomic traits (**Figure 4c)**.

### CASE STUDY

#### Dissection of the genetic architecture underlying flowering time variation in chickpea

Chickpea (*Cicer arietinum*) is a major food legume grown under semi-arid climatic conditions and is a vital source of dietary protein for the human population in developing countries. Further, due to growing awareness about the contribution of the meat industry to global warming, demands for plant-derived protein are also on the rise especially for most of the vegetarian human being in the developing world. To meet this soaring demand for high-quality plant-derived protein in global markets, protein-rich legumes like chickpea are expected to play a vital role. However, different abiotic stresses such as heat, drought, and end-of-season frost are known to shorten the growing season of chickpea, ultimately affecting seed yield in chickpea. Early flowering chickpea varieties which can escape such abiotic stresses are, therefore, vital to increase global chickpea production. Thus, understanding the genetic and molecular basis of flowering time in chickpea is of paramount importance. The efforts made so far to understand in this direction predominantly relied on bi-parental QTL mapping. These studies suggest that flowering time in chickpea is regulated by only a few major genes. These predominantly include Early flowering1 (*Efl1*), *Efl2*, *Efl3* and *Efl4* (**Lichtenzveig et al., 2006; Jamalabadi et al., 2013; Gaur et al., 2015; Weller and Ortega, 2015**). As bi-parental QTL mapping relies on genetic variation between only two parental accessions, it fails to provide a population-wide global genetic landscape of flowering and maturity time, the major yield contributing complex quantitative traits in chickpea. Thus, QTL mapping often misses many vital genes regulating traits of interest. GWAS is the proven strategy to overcome this bottleneck associated with conventional QTL mapping. Previously, an attempt has been made to conduct GWAS for flowering time traits in chickpea. This effort led to the identification of eight candidate genes, including an ortholog of Arabidopsis, PHOTOPERIOD-INDEPENDENT EARLY FLOWERING1 (PIE1) on chromosome 4 regulating flowering time chickpea (**Upadhyaya et al., 2015**). However, this study was based on only a smaller set of 92 diverse chickpea accessions and might have missed many vital genomic loci associated with flowering time due to the limited sample size. Thus, it will be interesting to conduct large-scale GWAS for flowering-related traits to uncover the global genetic architecture of flowering/maturity time in chickpea.

Therefore, in this study, to demonstrate the utility of GWAShub for large-scale GWAS using freely accessible genotype datasets, we decided to perform GWAS for diverse flowering/maturity-related traits utilizing publicly available genotype data for diverse chickpea accessions. For this, we phenotyped composite core panel consisting of 1954 diverse *desi* and *kabuli* chickpea accessions for two vital flowering-related traits, i.e. days to 50% flowering (DTF) and days to maturity (DTM). This panel is a subset of recently sequenced 3K chickpea accessions (**Varshney et al. 2021**) and the genotype information for these 3171 accessions is now available as part of GWAShub. The GWAS was then performed by uploading the phenotype data for both the DTF and DTM traits onto the GWAShub server. Visualization of phenotype data using GWAShub revealed wide-variation for both DTF and DTM traits measured in 1954 selected accessions (**Figure 5**). Although, both the traits were found to be normally distributed, to further improve the normality of phenotypes the Box-Cox transformation was applied to all studied traits (**Figure 5**). Using these normalized phenotypes, we further proceeded for GWAS utilizing CMLM and Blink algorithms, PCA covariate value of 2 and MAF of 0.05. The GWAS results were then assessed both independently and combinedly both for traits and algorithms using the visualization component of GWAShub. The results with both the algorithms were found to be identical thus we proceeded with Blink results for further analysis. This identified total of 10 different associations for DTF and about 11 different associations for DTM (**Figure 6**). Interestingly, the most prominent association for both DTF (Ca8:3934561; FDR p-Value 2.16E-32) and DTM (Ca8:3940395; FDR p-Value 8.11E-18) was not only found to be located on the same chromosome (Chromosome 8) but also found to coincide with the same haplotype block (LD_Block_CA8_3916685) (**Figure 7a, 8**). The haplotype block (LD_Block_CA8_3916685) spanned 195.159 kb (Chr8:3916685-4111843) and harbored a total of 24 genes, including the homolog of Arabidopsis CURLY LEAF (*CLF*) gene (*Ca_02166/LOC101514533*).

**Figure 5.**
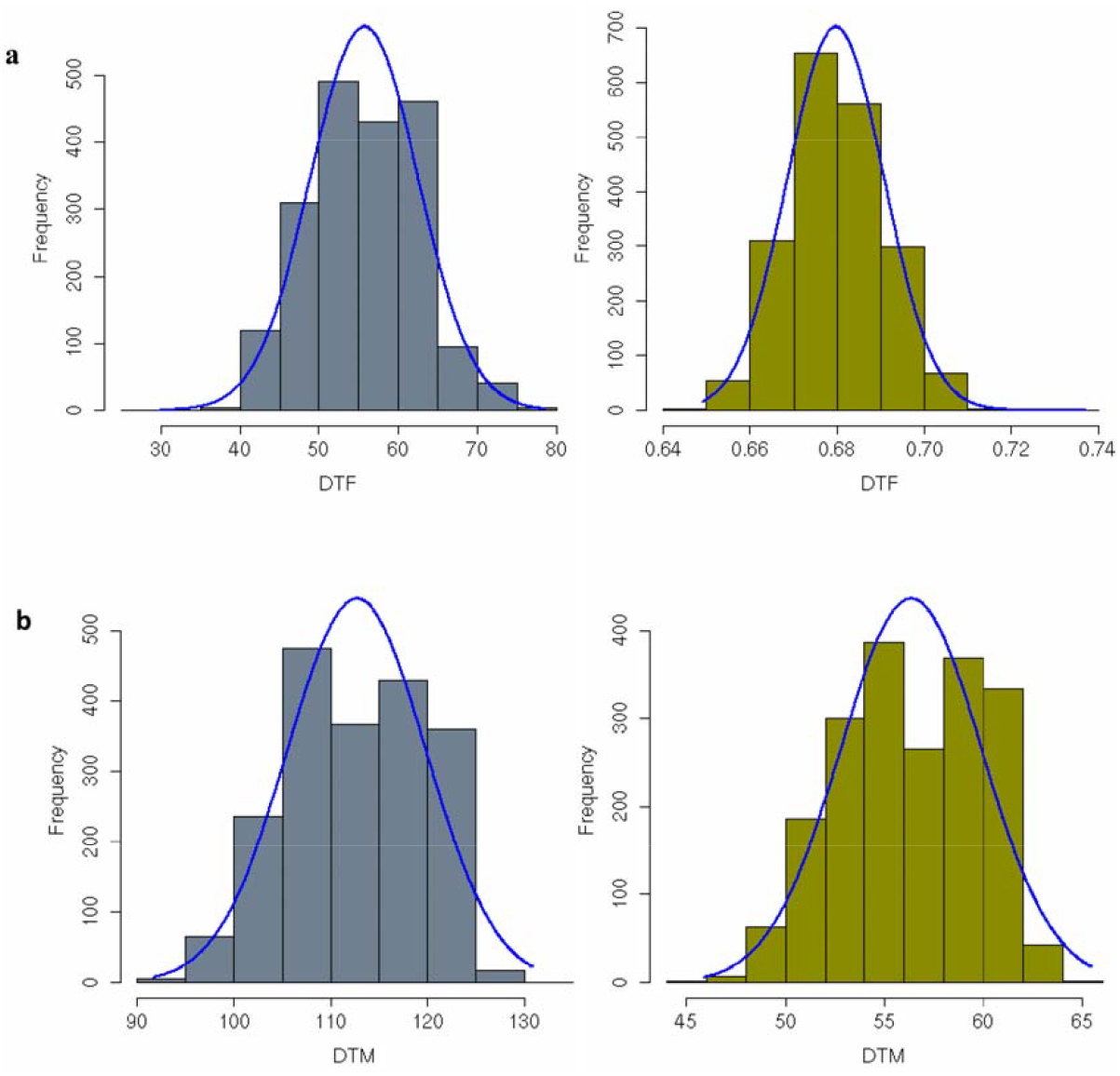
Diversity of flowering time related phenotypes in a composite core panel consisting of 1954 diverse chickpea accessions. **a)** Distribution of days to 50% flowering (DTF) phenotype before (grey) and after Box-Cox transformation (olive), and **b)** Distribution of days to 50% maturity (DTM) phenotype before (grey) and after Box-Cox transformation (olive). Frequency: number of chickpea accessions.

**Figure 6.**
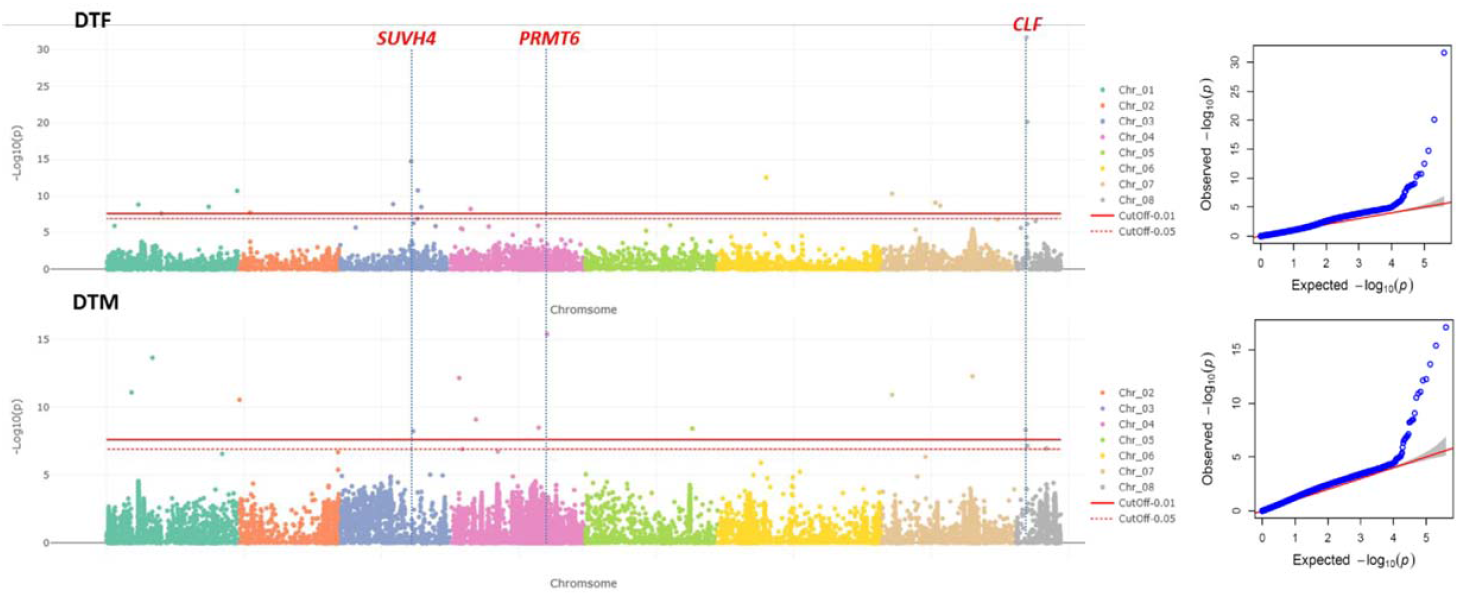
Manhattan plots and QQ-plots depicting results of genome wide association study (GWAS) for two flowering time traits i.e. days to fifty percent flowering (DTF) and days to maturity (DTM). The most probable candidate genes for associated loci arc marked on top in red. *SUVH4*: *SUPPRESSOR OF VARIEGATION 3–9 HOMOLOG;* PRMT6: PROTIEN HISTONE ARGNINE METHYL TRANSFERASE; *CLF: CURLY LEAF*

**Figure 7.**
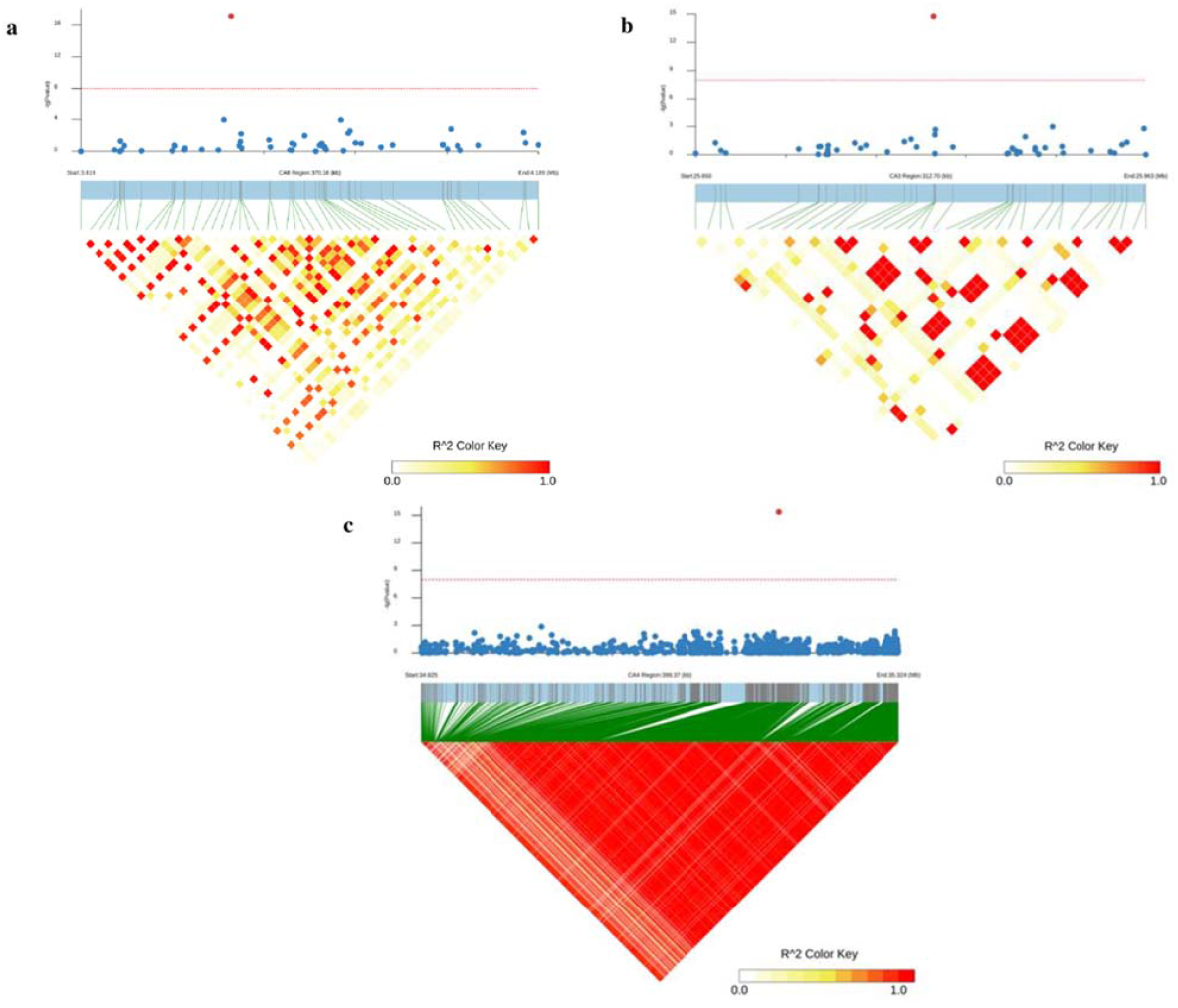
The LD heat-maps depicting haplotype blocks associated with different genomic loci regulating flowering time in chickpea. The haplotype blocks associated with **a)** chromosome 8 loci (Ca8:3916685-4111843 bp) detected for both day to 50% flowering (DTF) and days to maturity (DTM), **b)** chromosome 3 loci (Ca3:25747849 – 25870747 bp) regulating DTF and **c)** chromosome 4 loci (Ca4:35024272-35224272 bp) regulating DTM.

**Figure 8.**
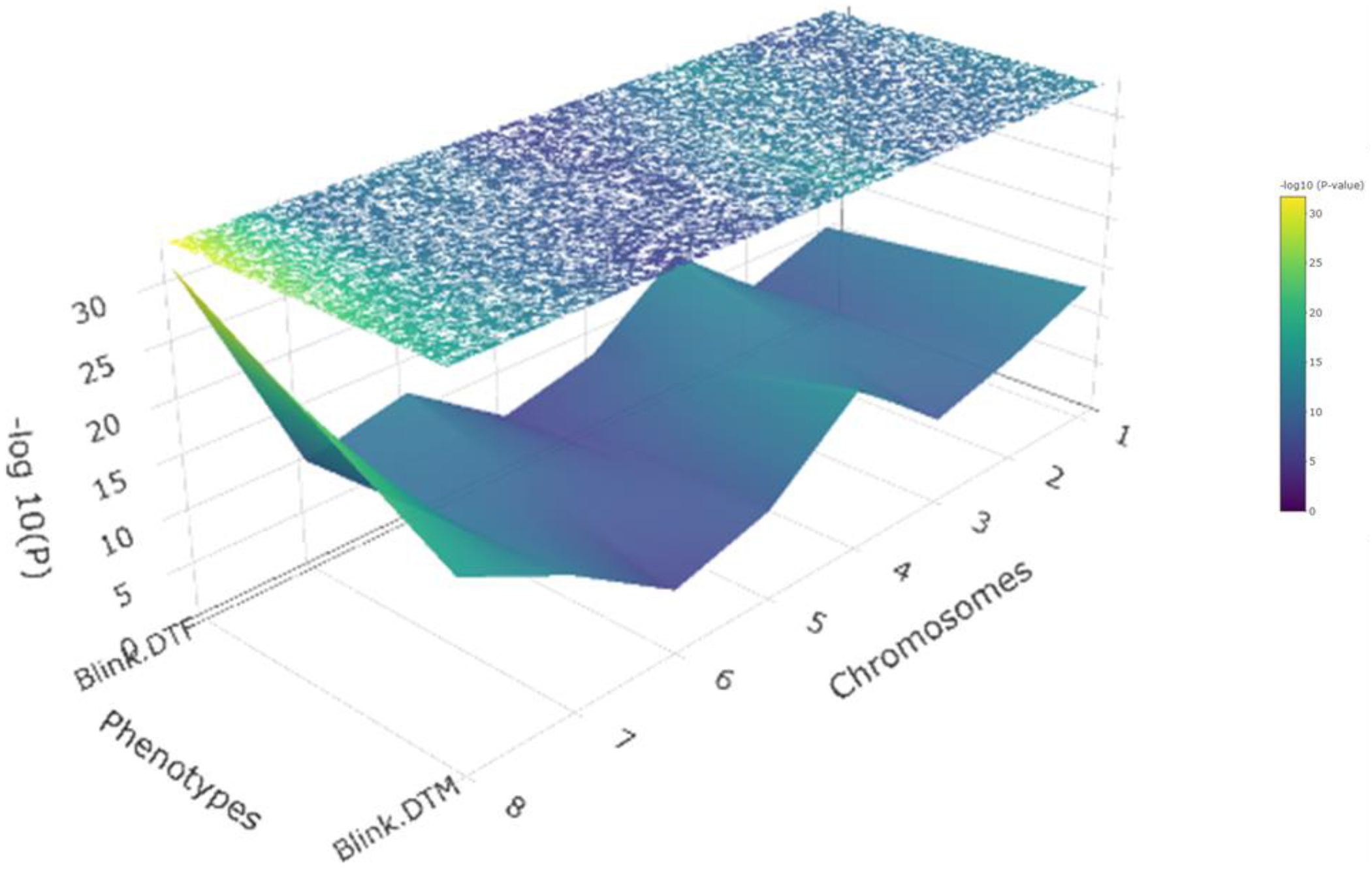
Exploration of pleiotropy at the genome-wide level using pheGWAS graph of two different flowering time-related phenotypes i.e., day to 50% flowering (DTF) and days to maturity (DTM). The highlighted region in the heatmap (elevated −log10 (p) values) suggests a pleiotropic loci on chromosome 8 controlling both DTF and DTM.

The CLF encodes for histone-lysine N-methyltransferase protein, belonging to the polycomb group (PcG) of proteins. The PcG proteins are known to form multisubunit protein complexes called polycomb repressive complex 1 (PRC1) and PRC2 which epigenetically regulate gene expression of target genes by methylation of histone tail. In Arabidopsis, CLF is known to be an important component of PRC2 and catalyzes the trimethylation of histone H3 lysine 27 (H3K27me3) (**Schubert et al. 2006**). Further, CLF is shown to have an essential role in maintaining epigenetic repression of FLOWERING LOCUS T (FT), which is a major gene involved in the transition from the vegetative to the reproductive phase. In addition to this, CLF has also been shown to be involved in epigenetic repression of flower homeotic genes like AGAMOUS (AG)/SHOOT MERISTEMLESS (STM) as a consequence of which loss of function CLF mutants flower early compared to wild type **(Goodrich et al. 1997; Schubert et al. 2006; Jiang et al. 2008; Lopez-Vernaza et al. 2012; Yan et al. 2018**). Further, the protein-protein interaction network analysis using the STRING database revealed interaction of CLF with many vital flowering related proteins including VERNALIZATION 2 (VRN2), RELATIVE OF EARLY FLOWERING 6 (REF6), EARLY FLOWERING 6 (ELF6), thus emphasizing the potential role of *CLF* in regulating flowering time in chickpea (**Figure 9, Gendall et al. 2001; Noh et al. 2004**). Besides, the recent study involving whole-genome sequencing of more than 3K wild and cultivated chickpea accessions revealed the genomic locus harboring *CLF* had undergone selection during domestication (**Varshney et al. 2021**). Thus, these multitudes of evidence suggest *CLF* as the most probable candidate gene underlying the major locus (Ca8:3934561; FDR p-Value 2.16E-32) regulating flowering time in chickpea. In contrast to this major locus on chromosome 8, all other detected loci were found to be associated with only one of the two traits.

**Figure 9.**
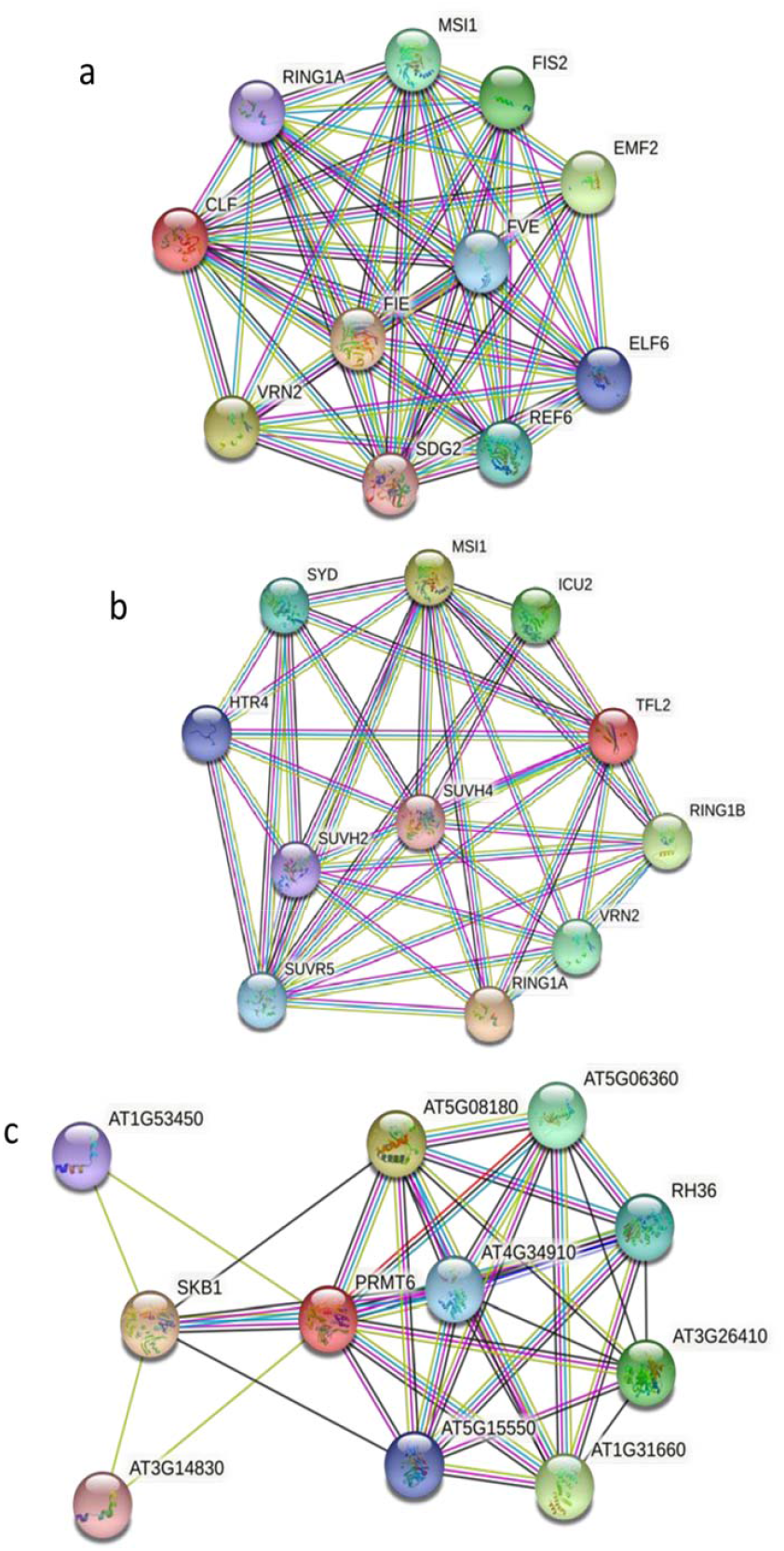
Protein-protein interaction networks of candidate genes underlying three major loci identified for two flowering time related traits i.e. day to 50% flowering (DTF) and days to maturity (DTM) using GWAS: a) CURLY LEAF (CLF), b) KRYPTONITE (KYP)/SUVH4 and c) protein arginine N-methyltransferase 6 (*PRMT6*).The color nodes represent query protein and first shell of interactors; Edges represent protein-protein associations (protein jointly contributing to same function), these include known protein interactions such as those curated databases (blue), experimentally determined interactions (violet). Similarly, the black and yellow edges represent the co-expression and co-occurrence in literature text, respectively.

A prominent association was identified for the DTF trait on chromosome 3 (Ca3:25747849; FDR p-Value 1.78E-15). The haplotype block associated with this locus (LD_Block_CA3_25747849) spanned 122.899 kb (Ca3: 25747849 – 25870747) and harbored only a single gene, KRYPTONITE (*KYP*)/SUVH4 (*Ca_23882/LOC101508428*), which encodes SET domain-containing H3K9 methyltransferase. The SET domain proteins are known to function in coordination with histone deacetylases (HDACs) to regulate flowering time in Arabidopsis (**Ning et al. 2019**). Further, in the case of the rice SET domain group (SDG) protein-encoding gene SDG712 which is responsible for H3K9me2 specific methylation is known to regulate flowering time in rice (**Zhang et al. 2021b**). Conversely, JmjC DOMAIN-CONTAINING PROTEIN 27 (JMJ27) which has an H3K9me1/2 demethylase activity regulates flowering time in Arabidopsis by negatively modulating the major flowering regulator CONSTANS (CO) and positively modulating FLOWERING LOCUS C (FLC) (**Dutta et al. 2017; Kinoshita and Richter, 2021**). These reports emphasize the crucial role of H3K9 methylation marks in the regulation of flowering-related genes in plants. Further, the STRING analysis revealed potential interaction of SUVH4 with well-known regulators of flowering time like TERMINAL FLOWER 2 (*TFL2*), *VRN2*, *SYD*, etc. (**Gendall et al. 2001; Nakagawa et al. 2002; Kotake et al. 2003**). Thus, SUVH4 can be studied further for its potential role in the regulation of DTF in chickpea. Similarly, a major locus on chromosome 4 (Ca4:35224030; FDR p-Value 1.78E-15) was detected for DTM. The haplotype block associated with this locus (LD_Block_CA4_35024272) spanned a genomic interval of ~ 200 kb (Ca4: 35024272-35224272) harboring a total of 10 protein-coding genes. One of these genes, (*Ca_20004*/LOC101489300) was again found to encode for protein arginine N-methyltransferase 6 (*PRMT6*). The *PRMTs* such as *PRMT5/SKB1* and *PRMT10* are known to regulate flowering time in plants by fine-tuning the expression of FLC, a major flowering repressor gene ( **Wang et al. 2007; Zhang et al. 2021a**). Recently, AtPRMT6 has been shown to interact physically with a known flowering regulator Nuclear Factors YC3 (NF-YC3), NF-YC9, and NF-YB3 (Hou et al. 2014; Siriwardana et al. 2016). Further, PRMT6 is found to function redundantly with its two homologues PRMT4a and PRMT4b to regulate flowering time in *Arabidopsis* potentially by suppressing the expression of *FLOWERING LOCUS C*, which in turn suppresses *FLOWERING LOCUS T* (*FT*) (**Zhang et al. 2021a**). Thus, taking the aforementioned evidence into account, *PRMT6* identified in this study can be considered a potential candidate gene for regulating DTM in chickpea.

The major role of epigenetic modifications, like histone methylation and acetylation in the regulation of flowering time is now well established, especially in the model plant Arabidopsis. Interestingly, all the three candidate genes identified in this study, encode different proteins with histone methyltransferases activity. The histone methyltransferases act by modifying chromosomal histones which in turn leads to suppression/activation of target flowering-related genes. Thus, the epigenetic regulators seem to be the major contributors to flowering time variation existing in natural chickpea accessions. The functional studies aimed at deciphering the role of these identified epigenetic regulators are expected to give further insights into the complex genetic architecture underlying flowering and maturity time in chickpea.

### SUMMARY AND FUTURE DIRECTIONS

GWAS has become an indispensable tool in the arsenal of contemporary plant geneticists for understanding the genetic architecture of complex plant traits. The recent availability of large-scale population-level sequence variation data for almost all major field and vegetable crops can further boost GWAS in crop plants. However, this requires access to expensive computational infrastructure and advanced technical skills essential to handle big biological datasets. Thus, the efficient utilization of publicly available sequence variation data remains challenging for resource-scarce plant researchers, especially from the developing world. Here, with the GWAShub server, we have attempted to address both of these critical challenges by providing a user-friendly interface to readily perform GWAS using public genotype datasets of diverse crops. The GWAS-hub not only provides access to curated genotype datasets but also allows easy-to-use implementation of well-structured GWAS workflow built using a variety of widely-tested popular computational tools to generate reproducible GWAS results. Further, multiple visualization options, extensive structural-functional annotation available in GWAShub allow easy visualization and rapid interpretation of GWAS results. Finally, the postGWAS functionality leverages the power of machine learning to prioritize the causal gene from the identified QTLs. Thus, we believe GWAShub will play an important role in making the wealth of publicly available genomic information readily accessible for GWAS and empower plant researchers to unravel the genetic architecture of complex quantitative traits in a self-reliant manner.

In future, we plan to extend support for all the major food crop plants for which population-level sequence variation data is available publicly. We are also working on adding extensive functional annotation including Gene Ontology (GO), Pfam protein families domains and KEGG pathway annotation to provide added convenience for the user while inferring genes and molecular pathways regulating trait of interest. Finally, to keep up with the latest advancements in GWAS in the pan-genomics era, we are working on adding functionality to conduct k-mer based association analysis to GWAShub. This will become available as a feature in the upcoming version.

## DATA AVAILABILITY

The complete results flowering time GWAS in chickpea can be accessed using the following link: http://gwashub.com/chickpeaGWAS/tmp/rishi543210_97428/processing/ALL_RES_1.php

## ACKNOWLEDGMENTS

Financial support for this study was provided by a research grant from the Department of Biotechnology (DBT), Government of India (102/IFD/SAN/2161/2013-14). AD acknowledges the DBT, India for research fellowship award.

